# Delta glutamate receptor conductance drives excitation of dorsal raphe neurons

**DOI:** 10.1101/855353

**Authors:** Stephanie C. Gantz, Khaled Moussawi, Holly S. Hake

## Abstract

The delta glutamate receptors, GluD1_R_ and GluD2_R_, are mysterious members of the ionotropic glutamate receptor family in that they are not gated by glutamate^1,2^. One theory is that they are scaffolding proteins or synaptic organizers strictly, rather than ion conducting channels. Although mutant forms and wild type channels have been reported to conduct^3,4,5^, conduction, gating, and biophysical properties of native GluD1_R_ remain unexplored. Here we show that the inward current induced by activation of α1-adrenergic receptors (α1-A_R_s) in the dorsal raphe nucleus (DR) is mediated by GluD1_R_. Native GluD1_R_ channels are functional ion channels that are constitutively active under basal conditions and α1-A_R_s increase the tonic current. This inward current is responsible for the α1-A_R_-dependent induction of persistent pacemaker-type firing of neurons in the DR. Given the extensive distribution of these receptors, the ionotropic nature of GluD_R_ is proposed to be widespread in the nervous system.

---

The dorsal raphe nucleus (DR) is the largest serotonergic nucleus in the brain and the predominant source of central serotonin (5-HT). Activation of Gα_q/11_ protein-coupled α1-adrenergic receptors (α1-A_R_s) in the DR produces a slow and long-lasting membrane depolarization^6^. *In vivo*, tonic noradrenergic input that activates α1-A_R_s is required for 5-HT neurons to fire action potentials^7,8^. Despite having a crucial role in regulating 5-HT neuron excitability, the ion channels responsible for the depolarization remain unknown.

## Activation of α1-A_R_ produces an EPSC

Electrophysiological recordings were made from DR neurons in acute brain slices from wild type mice at 35° C in the presence of NMDA_R_, AMPA_R_, Kainate_R_, GABA-A_R_, and 5-HT1A_R_ antagonists. With cell-attached recordings, a train of 5 electrical stimuli (60 Hz) produced firing in previously quiescent neurons, which was blocked by application of the α1-A_R_ antagonist, prazosin (100 nM, Fig. 1a). The excitation produced 20±5 action potentials (APs) that lasted 9.0±3.0 s, with a latency of 650.6±0.1 ms from onset of stimulation to the first AP (Fig. 1b-e). In whole-cell recording using a potassium-based internal solution, the same train of electrical stimuli produced prolonged AP firing (Fig. 1f). In voltage-clamp mode (V_hold_ −65 mV), the same stimulation produced a slow and long-lasting (27.4±2.3 s, n=10) excitatory postsynaptic current (EPSC, Fig. 1f) that was eliminated by application of prazosin (Fig. 1g). Prazosin had no effect on basal whole-cell current (−3.8±3.4 pA, p=0.232, n=10, not shown) indicating a lack of persistent inward current due to noradrenaline (NA) tone. Application of tetrodotoxin (1 μM) reversibly abolished the α1-A_R_-EPSC, demonstrating a dependence on presynaptic action potentials (Fig. 1h). Disruption of the vesicular monoamine transporter with reserpine (1 μM) or removal of external Ca^2+^ also eliminated the α1-A_R_-EPSC, indicating NA release is vesicular (Fig. 1i,j). To test whether α1-A_R_-EPSCs were dependent on G protein-signaling, recordings were made with an internal solution containing GDP-βS-Li_3_ (1.8 mM) instead of GTP. Disruption of G protein signaling with intracellular dialysis of GDP-βS-Li_3_ eliminated the α1-A_R_-EPSC within 5-20 min post-break-in (p=0.004, n=9), whereas dialysis with LiCl alone had no effect on the α1-A_R_-EPSC amplitude (p=0.625, n=4, Fig. S1a). These findings demonstrate a cell-autonomous requirement of G protein signaling in the generation of the α1-A_R_-EPSC.

**Figure 1.**
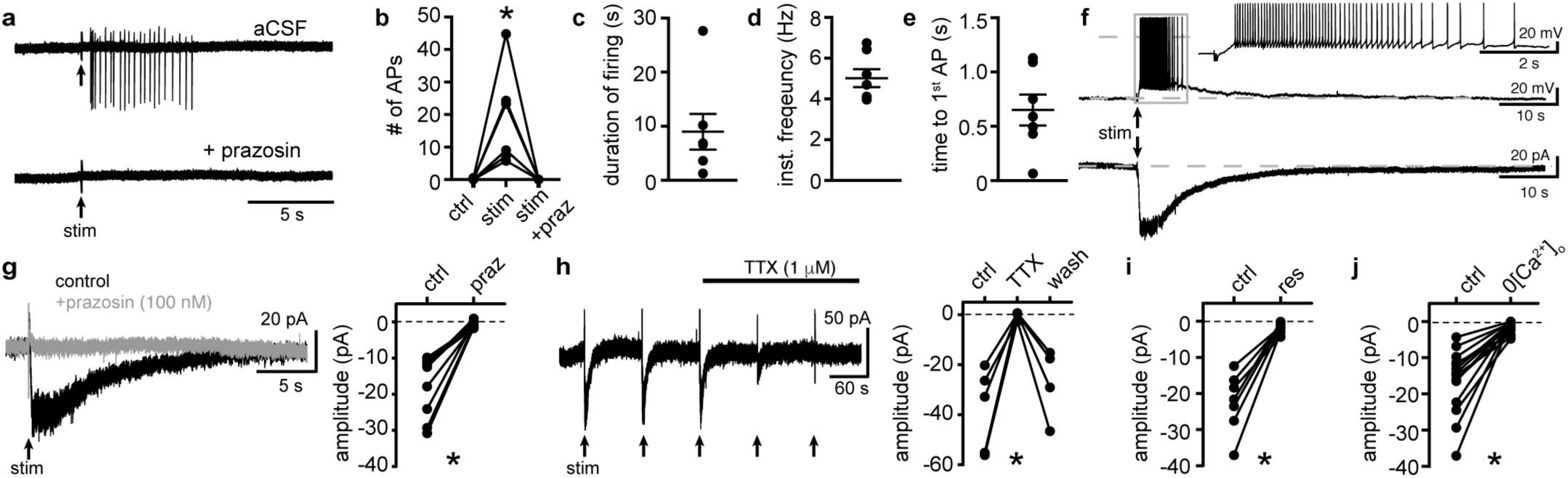
Electrical stimulation evokes long-lasting AP firing produced by an α1-adrenergic receptor-dependent EPSC. **a**, Representative traces of cell-attached recording where stimulation of the brain slice (5 stims at 60 Hz) produced AP firing that was abolished by application of the α1-adrenergic receptor antagonist, prazosin (100 nM). **b**, Plot of number of APs showing the stimulation-induced increase in frequency (p=0.004, n=6). **c**, Plot of duration of AP firing. **d**, Plot of mean instantaneous frequency of AP firing over the first 10s of firing. **e**, Plot of the latency from stimulation onset to the first AP. **f**, Example whole-cell recordings in the same cell, where electrical stimulation of the slice produced prolonged AP firing in current-clamp (upper trace) and a slow EPSC in voltage-clamp mode (lower trace). **g**, Application of prazosin eliminated the slow EPSC shown in a representative trace (left, baseline adjusted) and in grouped data (right, p=0.002, n=10). **h**, Application of tetrodotoxin (TTX, 1 μM) reversibly eliminated the α1-A_R_-EPSC shown in a representative trace (left, α1-A_R_-EPSC evoked every 90s (arrows)) and in grouped data (right, p=0.009, n=4). **i**, Plot of the inhibition of α1-A_R_-EPSC amplitude by application of reserpine (res, 1 μM, p=0.016, n=7). **j**, Plot of the inhibition of α1-A_R_-EPSC amplitude by removal of external Ca^2+^ (0[Ca^2+^]_o_, p=0.0001, n=14). Line and error bars represent mean±SEM, * denotes statistical significance.

## Biophysical properties of the channel

Under our recording conditions, membrane resistance (R_m_) decreased during the α1-A_R_-EPSC, indicative of ion channels opening (Fig. 2a). Membrane noise variance (σ^2^) increased during the EPSC compared to membrane noise under basal conditions (Fig. S2a,b). The α1-A_R_-EPSC σ2-amplitude relationship was well-fit by linear regression, yielding a measurement of −1.16 pA unitary current (Fig. S2c). Voltage ramps from −120 to −10 mV (1 mV/10 ms) before and during the α1-A_R_-EPSC (Fig. 2b) showed that the current reversed polarity at −28.6±2.4 mV (Fig. 2b-d). Exogenous application of NA (30 μM) produced an inward current (I_NA_) with a similar reversal potential (E_rev_, - 25.1±2.9 mV, Fig. 2d). Replacing 126 mM [Na^+^]_o_ with N-methyl D-glucamine (NMDG) completely abolished inward I_NA_, suggesting Na^+^ as the prominent charge carrier (Fig. 2e). Increasing [K^+^]_o_ from 2.5 to 6.5 or 10.5 mM, expected to shift E_K_ from −107 to −81 and −69 mV, respectively, had no effect on the α1-A_R_-EPSC amplitude at V_hold_ −65 mV nor −120 mV, but produced a depolarizing shift in E_rev_ of the α1-A_R_-EPSC (Fig. S3a-c), suggesting the channel is also permeable to K^+^. Removal of external MgCl_2_ had no significant effect on E_rev_, nor on the amplitude of I_NA_ (Fig. S4a,b). Removal of external CaCl_2_ also had no significant effect on E_rev_, but augmented inward I_NA_, (Fig. S4a,b). Taken together, the data suggest that α1-A_R_-dependent current, whether produced by vesicular NA release or exogenous NA is carried through a mixed cation channel, with inward current carried predominantly by Na^+^ entry.

**Figure 2.**
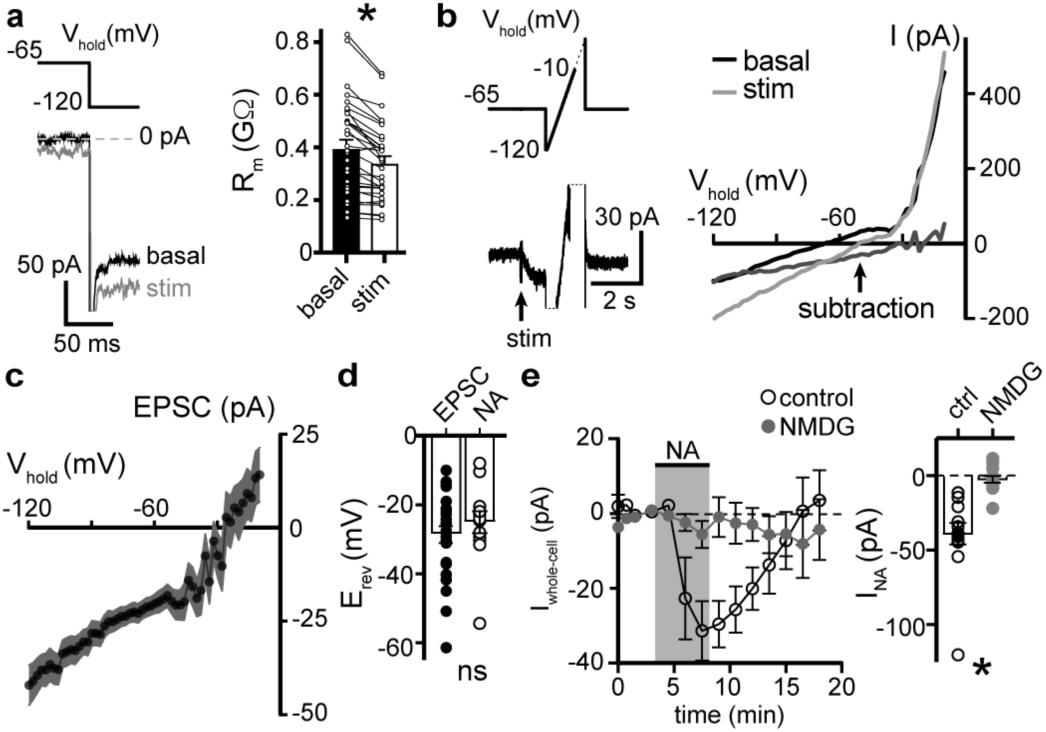
α1-adrenergic receptor-dependent inward current is carried by sodium entry. **a**, Membrane resistance (R_m_, ΔV −65 to −120 mV) decreased during the α1-A_R_-EPSC indicating an opening of ion channels, as shown in an example trace (left) and in grouped data (right, p<0.0001, n=31). **b**, Representative trace of current-voltage relationships of the α1-A_R_-EPSC (subtraction), determined by subtracting current measured during slow voltage ramps (1 mV/10 ms, analyzed from −120 to −10 mV) at the peak of the α1-A_R_-EPSC (stim) from current measured in control conditions just prior to stimulation (basal). Current generated during ramps were truncated for clarity. **c**, Current-voltage relationship of the α1-A_R_-EPSC from grouped data. **d**, Plot of reversal potentials (E_rev_) of the α1-A_R_-EPSC and I_NA_ (p>0.999, n=26&14). **e**, Replacing 126 mM NaCl with NMDG eliminated inward I_NA_, shown in a time-course plot (left) and grouped data (right, V_hold_ −65 mV, p<0.0001, n=14&13). Line and error bars/shaded area represent mean±SEM, * denotes statistical significance, ns denotes not significant.

Measurements of E_rev_ assume voltage-independence of the channel and the signaling mechanism between α1-A_R_ and the channel. To test for voltage-dependence, we employed a two-pulse voltage-step protocol. Current was measured at V_hold_ −120 mV following a conditioning pre-pulse (−120 to 30 mV, 150 ms) before and after application of NA (Fig. S2f,g). I_NA_ was isolated by subtracting the current under basal conditions from the current during NA. Conductance (G_NA_) was calculated, using an E_rev_ of −25.1 mV. Conditioning depolarizing pre-pulses incrementally reduced G_NA_ and the increase in membrane noise induced by NA measured at V_hold_ −120 mV (Fig. S2h-i), demonstrating voltage-dependence of inward I_NA_, such that prior depolarization reduced conductance.

## α1-A_R_s modulate tonic GluD1_R_ current

Activation of Gα_q/11_ protein-coupled receptors (G_q_PCRs), namely metabotropic glutamate mGlu_R_s, muscarinic acetylcholine M1 (mACh_R_s), or α1-A_R_s produces slow, noisy inward currents throughout the central and peripheral nervous system. Multiple mechanisms have been reported to underlie the inward current including: inhibition of K^+^ current (including leak, Ca^2+^-activated, and Kv7/M-current)^9–12^, modulation of TTX-sensitive persistent Na^+^ current^13^, and activation of transient potential receptor canonical (TRPC)^14,15^, Na^+^-leak (NALCN)^16^, or GluD_R_ channels^4,5^. GluD1_R_-encoding (Grid1) mRNA is expressed widely throughout the brain, with notably high levels in the DR^17,18^. To assess involvement of GluD1_R_, we applied a channel-selective toxin. Akin to other Ca^2+^-permeable ionotropic glutamate receptors, GluD_R_ channels are blocked by the Joro spider toxin, 1-Naphthyl acetyl spermine (NASPM)^5,19^. Application of NASPM (100 μM, 6 min) blocked the α1-A_R_-EPSC (96.0±12.5% reduction), which recovered to baseline after >30 min of wash (Fig. 3a,b,d,f). NASPM alone produced an apparent outward current (I_NSP_) of 20.5±3.7 pA with an E_rev_ of −31.4±4.8 mV (Fig. 3a,c,d; Fig. S5) and a reduction in membrane noise (Fig. S2d,e). Upon washout, I_NSP_ reversed with a similar time course of recovery of the α1-A_R_-EPSC (Fig. 3d). I_NSP_ was associated with an increase in R_m_ (Fig. 3e) indicating a closure of channels. Replacing 126 mM [Na^+^]_o_ with NMDG eliminated I_NSP_ (Fig. S5a). Thus, I_NSP_ was due to block of tonic Na^+^-dependent inward current. I_NSP_ was not dependent on prior electrical stimulation of the brain slice, as the magnitude of I_NSP_ was similar between stimulated and unstimulated brain slices (Fig. S5b,c). Given that NASPM is an open-channel blocker^20^, we tested whether electrically evoking an α1-A_R_-EPSC during application of NASPM was required for block. After obtaining a steady α1-A_R_-EPSC baseline, NASPM was applied for 6 min without stimulating the brain slice. The α1-A_R_-EPSC was blocked when stimulation was reapplied (Fig. 3f), indicating that the channels underlying the α1-A_R_-EPSC were blocked. Thus, the α1-A_R_-EPSC is carried by channels that are at least transiently open at rest and may be the same channels that underlie the NASPM-sensitive tonic inward current.

**Figure 3.**
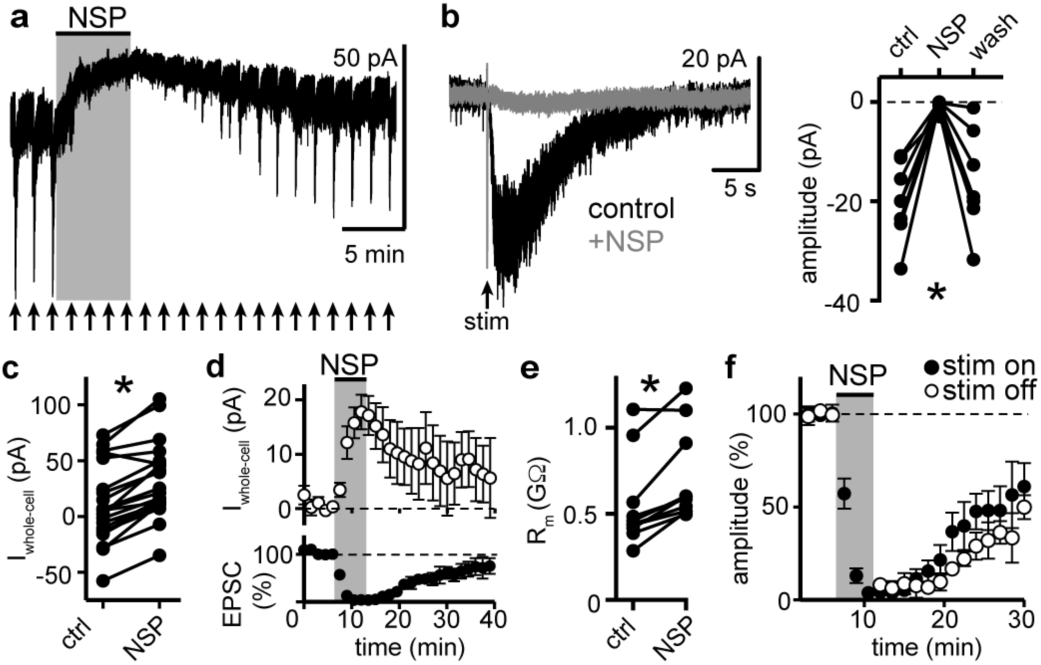
NASPM blocks the α1-A_R_-EPSC and a tonic inward current. **a**, Example whole-cell voltage-clamp recording of the basal whole-cell current and the α1-A_R_-EPSC evoked every 90s (arrows) prior to, during, and after application of NASPM (NSP, 100 μM). **b**, NASPM completely eliminated the α1-A_R_-EPSC shown in representative traces (left, baseline adjusted) and in grouped data (right, p=0.002, n=10). **c**, Application of NASPM produced an apparent outward current (p<0.0001, n=21). **d**, Time course of the inhibition of the α1-A_R_-EPSC amplitude (bottom) and apparent outward current (top) by application of NASPM. **e**, Membrane resistance (R_m_, ΔV −65 to −75 mV) increased during application of NASPM, indicating the apparent outward current was due to ion channels closing (p=0.004, n=10). **f**, Time course of the inhibition of the α1-A_R_-EPSC amplitude by application of NASPM, demonstrating identical block of the α1-A_R_-EPSC whether or not α1-A_R_-EPSCs were evoked during NASPM application. Line and error bars represent mean±SEM, * denotes statistical significance.

GluD_R_s bind the amino acids D-serine and glycine which inhibit constitutively open mutant GluD_R_ current^21,22^. Application of D-serine (10 mM, 13.5 min) reduced the α1-A_R_-EPSC amplitude by 49.7±9.6% (Fig. S6a,c). Application of glycine (10 mM, 4.5 min) reduced the α1-A_R_-EPSC amplitude by 70.9±11.0% (Fig. S6b,c). Lastly, application of the glutamate receptor antagonist kynurenic acid (1 mM, 10.5 min) reduced the α1-A_R_-EPSC amplitude by 65.6±8.3% (Fig. S6c).

Next, a viral genetic strategy was used to functionally delete GluD1_R_ by targeting the encoding gene, *GRID1*, via CRISPR/Cas9 (Fig. S7a,b). In brief, a cocktail of AAV1 viruses encoding Cas9, and mouse Grid1 guide RNA with enhanced green fluorescent protein (eGFP) reporter (GRID1) was microinjected into the DR of wild type mice. A separate cohort received microinjections of a cocktail of AAV1 viruses encoding Cas9 and eGFP reporter (control). Brain slices were prepared after >4 weeks and the DR was microdissected and frozen on dry ice to assess the mutation of *GRID1*. Restriction enzyme site-digested PCR confirmed *in vivo* mutation of *GRID1* at the expected site (Fig. S7c). In a separate cohort, brain slices were prepared and whole-cell voltage-clamp recordings were made from infected and uninfected neurons visualized in brain slices by expression of eGFP. In eGFP^+^ neurons from control mice, electrical stimulation produced a decrease in R_m_ and an α1-A_R_-EPSC, and application of NA caused inward I_NA_ (Fig. 4). However, in eGFP^+^ neurons from GRID1 mice, electrical stimulation did not change R_m_ (Fig. 4a) and no α1-A_R_-EPSC was detected above baseline noise (Fig. 4b). In addition, inward I_NA_ was substantially smaller in eGFP^+^ neurons from GRID1 mice, as compared to eGFP^+^ neurons from control mice (Fig. 4c). In the same slices from GRID1 mice, eGFP-neurons still had an α1-A_R_-EPSC and inward I_NA_ (Fig. 4a,c). Lastly, application of NASPM produced an apparent outward current in eGFP^+^ neurons from control mice, but not from GRID1 mice (Fig. 4d). Taken together, these results demonstrate that conduction through GluD1_R_ is necessary for the α1-A_R_-EPSC and the NASPM-sensitive tonic inward current.

**Figure 4.**
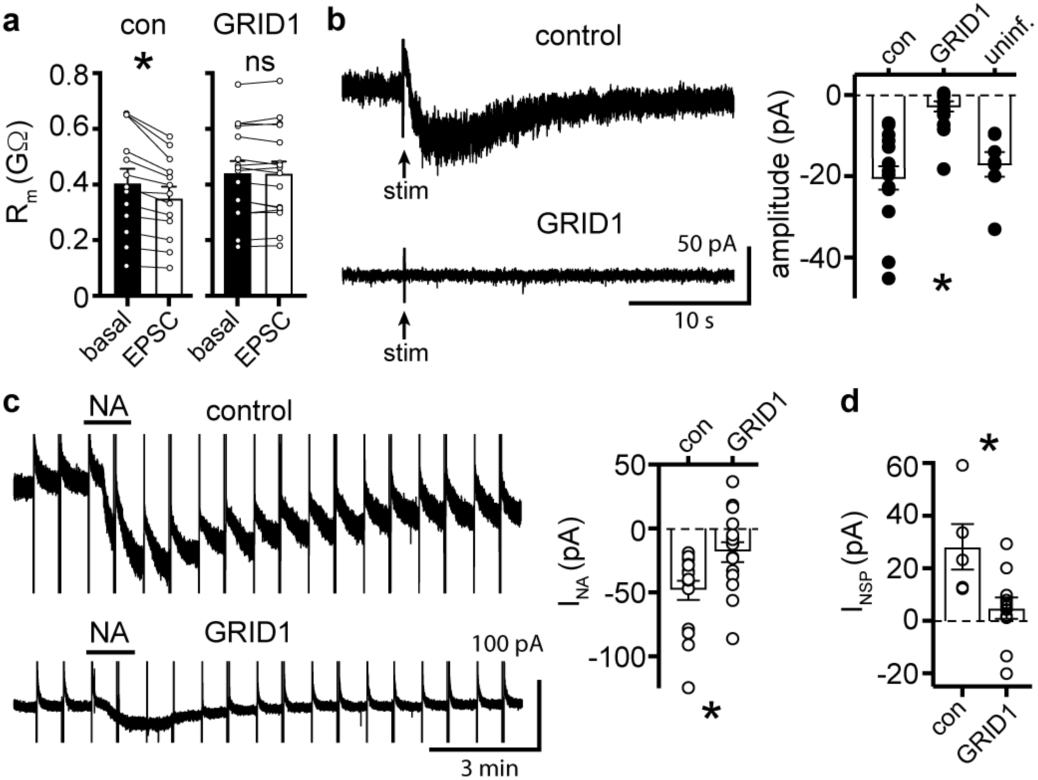
The α1-A_R_-EPSC is eliminated by targeting of GluD1 receptors via CRISPR/Cas9. **a**, Membrane resistance (R_m_, ΔV −65 to −120 mV) decreased after stimulation in infected neurons from mice injected with AAV-Cas9 and AAV-empty (control, p=0.0002, n=13), but not in infected neurons from mice injected with AAV-Cas9 and AAV-GRID1 (GRID1, p=0.562, n=16). **b**, Representative traces (left) and grouped data (right, p<0.0001, n=16&16) demonstrating the presence of an α1-A_R_-EPSC in infected neurons from control mice, but not from GRID1 mice. Neighboring uninfected neurons from GRID1 mice (uninf.) had an α1-A_R_-EPSC that was indistinguishable from infected neurons from control mice (p>0.999, n=7). **c**, Targeting GluD1_R_ reduced the inward current to NA (I_NA_, 30 μM) as compared to infected neurons from control mice, as shown in representative traces (left) and grouped data (right, p=0.009, n=16&16). Capacitive transients and ramp-induced currents have been truncated for clarity. **d**, Targeting GluD1_R_ reduced the tonic inward current revealed by application of NASPM (100 μM, I_NSP_) as compared to infected neurons in control mice (p=0.009, n=5&11). Line and error bars represent mean±SEM, * denotes statistical significance, ns denotes not significant.

## Relevance of GluD1_R_ to the dorsal raphe

*In vivo*, 5-HT neurons in the DR require NA release and subsequent activation of α1-A_R_s to maintain AP firing^8^. Activation of α1-A_R_s in the DR by exogenous agonist was reported to depolarize neurons through net reduction of K^+^ conductance and by activation of an unidentified non-K^+^ conductance^23^. More recently, Brown *et al*., reported that α1-A_R_s activation in the DR induces Na^+^-dependent inward current with an E_rev_ of −23 mV^24^, similar to our findings. Our study identifies GluD1_R_ as the channel that carries this mixed cation current, indicating that modulation of GluD1_R_ is a key constituent in driving persistent AP firing of the 5-HT neurons. In principle, inward GluD1_R_ current may bring the membrane potential to threshold, but recruitment of other voltage-gated ion channels is expected to underlie the persistent pacemaker-like activity.

## Ionotropic nature of GluD1_R_

GluD_R_ have been characterized as synaptic organizers, regulating LTD, endocytosis and trafficking of AMPA_R_, formation of excitatory and inhibitory synapses, and spine density^3,25–27^, without the requirement of ion conduction. Similarly, NMDA_R_ signal through non-ionotropic or “metabotropic” mechanisms where ion conduction is not required, to regulate LTD, AMPA_R_ endocytosis, and spine morphology^28^. The ionotropic nature of GluD_R_ does not conflict with its known role as a scaffold protein, but rather expands the similarities between NMDA_R_ and GluD_R_.

The largest obstacle towards understanding the ionotropic nature of GluD_R_ is the lack of known agonist and inability to gate the intact channel. The majority of studies have been performed on constitutively open mutant or chimeric channels^29,30^. Two prior studies have demonstrated that in heterologous systems and brain slices, activation of metabotropic glutamate receptors (mGlu_R_) produces an inward current carried by GluD1_R_^5^ or GluD2_R_^4^, concluding mGlu_R_ activation triggers rather than modulates gating of GluD_R_. The congruous explanation of our results is that, in DR neurons, GluD1_R_ are functional and open under basal conditions, carrying subthreshold, tonic Na^+^ current. Activation of α1-A_R_s, by exogenous agonist or synaptic NA release, excites DR neurons by increasing GluD1_R_ current.

In general, the kinetics of iGlu_R_ synaptic currents are controlled by the lifetime of the receptor-agonist complex and the rates of desensitization and deactivation. In brain slices, NMDA_R_ carry tonic inward current^31^ attributed to slow desensitization and ambient levels of glutamate and glycine. What remains to be understood are the conditions that permit GluD1_R_ opening, and why their activation has been largely elusive in heterologous expression systems. Reminiscent of times before the discovery of glycine as a necessary co-agonist at NMDA_R_^32,33^, it may be that an endogenous agonist needed for gating is present in brain slices. Alternatively, it is possible GluD1_R_ are gated by an intracellular factor or require expression of accessory or interacting proteins^34^. Tonic activation of α1-A_R_s cannot explain the tonic inward current as α1-A_R_ antagonism did not change basal whole-cell current.

The mechanism by which α1-A_R_s increase GluD1_R_ current also remains to be described and may be distinct from tonic activation. It is well-established that GqPCRs, especially mGlu_R_ and mACh_R_, bidirectionally change NMDA_R_ and AMPA_R_ currents, producing the two major forms of synaptic plasticity, long-term potentiation (LTP) and long-term depression (LTD), in part through a variety of postsynaptic mechanisms^35^. To our knowledge, the duration of the α1-A_R_-EPSC (~27 s) is exceptional for any synaptic current and more closely resembles the duration of short-term synaptic plasticity; for instance, endocannabinoid-mediated short-term depression^36^. G_q_PCRs can activate phospholipase C which hydrolyzes the integral membrane lipid, phosphatidylinositol 4,5-bisphosphate (PIP2). PIP2 stabilizes Kv7 channels such that PIP2 hydrolysis following mACh_R_ activation accounts for inhibition of M-current^37^. By the same signaling cascade, G_q_PCRs stimulate the production of endocannabinoids that can act directly on membrane ion channels^38^. Thus, one possibility is that α1-A_R_s modulate GluD1_R_ through membrane lipid signaling. Largely, it remains to be seen whether the intracellular signaling cascades known to affect NMDA_R_ and AMPA_R_ currents generalize to GluD_R_.

Using heterologous systems or constitutively open mutants, GluD_R_ current reverses polarity around 0 mV^39^, akin to AMPA_R_ and NMDA_R_, while our results show an E_rev_ of ~ −30 mV. The difference may be due space-clamp error in brain slices, since the magnitude of subtracted current is small relative to total current at depolarized potentials. Activation of α1-A_R_s in the DR also inhibits A-type K^+^ current^40^. Thus, E_rev_ measurements may include inhibition of a voltage-gated K^+^ conductance that is not contributing directly to the α1-A_R_-EPSC but is activated during voltage ramps at V_hold_ >-65 mV. Tail current analysis revealed voltage-dependence of I_NA_, such that prior depolarization reduced conductance. These data may reflect block of GluD1_R_ by endogenous intracellular polyamines, as established for calcium-permeable AMPA_R_ and Kainate_R_^41^. Another important consideration is that our measurements may be subject to voltage-dependence of the signaling pathway between α1-A_R_s and GluD1_R_s. There are reports of inward currents generated by GqPCR activation with E_rev_s between −40 and −23 mV^13,24,42,^ measured under different recording conditions, suggesting a commonality that is unlikely to be due to space-clamp error alone. Taken together, measurements here should be considered an approximate estimate of GluD1_R_, and more precisely as the current-voltage relationship of the α1-A_R_-GluD1_R_ signaling complex.

## Summary

Native GluD1_R_ are functional ion channels that carry ionic current. GluD1_R_ underlie the α1-A_R_-mediated depolarization of DR neurons that drives AP firing *in vivo*. Many of the biophysical properties of the GluD1_R_ channel are shared with other members of the ionotropic glutamate receptor family. This study lays the foundation to investigate the ionotropic role of GluD1_R_ in excitatory synaptic transmission and neuronal excitability, expanding upon the wealth of knowledge of pharmacology and regulatory elements established for NMDA_R_ and AMPA_R_ signaling.

## METHODS

### Animals

All studies were conducted in accordance with the National Institutes of Health Guide for the Care and Use of Laboratory animals with the approval of the National Institute on Drug Abuse Animal Care and Use Committee. Wild type C57BL/6J (>3 months old) mice of either sex were used. Mice were group-housed on a 12:12 h reverse light cycle.

### Brain slice preparation and electrophysiological recordings

Mice were deeply anesthetized with isoflurane and killed by decapitation. Brains were removed quickly and placed in warmed modified Krebs’ buffer containing (in mM): 126 NaCl, 2.5 KCl, 1.2 MgCl_2_, 1.2 CaCl_2_, 1.2 NaH_2_PO_4_, 21.5 NaHCO_3_, and 11 D-glucose with 5 μM MK-801 to reduce excitotoxicity and increase viability, bubbled with 95/5% O_2_/CO_2_. In the same solution, coronal DR slices (220 μm) were obtained using a vibrating microtome (Leica 1220S) and incubated at 32 °C >30 minutes prior to recording.

Electrophysiological recordings were made at 35 °C with a Multiclamp 700B amplifier (Molecular Devices), Digidata 1440A A/D converter (Molecular Devices), and Clampex 10.4 software (Molecular Devices) with borosilicate glass electrodes (King Precision Glass) wrapped with Parafilm to reduce pipette capacitance. Pipette resistances were 1.8-2.8 MΩ when filled with an internal solution containing, (in mM) 104.56 K-methylsulfate, 5.30 NaCl, 4.06 MgCl_2_, 4.06 CaCl_2_, 7.07 HEPES (K), 3.25 BAPTA(K4), 0.26 GTP (sodium salt), 4.87 ATP (sodium salt), 4.59 creatine phosphate (sodium salt), pH 7.32 with KOH, mOsm ~285, for whole-cell patch-clamp recordings. Series resistance was monitored throughout the experiment. Cell-attached recordings were made from quiescent neurons in slice, using pipettes filled with modified Krebs’ buffer. Reported voltages are corrected for a liquid junction potential of −8 mV between the internal solution and external solution. All drugs were applied by bath application. All experiments were conducted with NMDA_R_, AMPA_R_, Kainate_R_, GABA-A_R_, and 5-HT1A_R_ antagonists in the external solution. In addition, a α2-adrenergic receptor antagonist was added for experiments where NA was applied and a glycine receptor antagonist was added when glycine was applied. Unitary current was calculated from fluctuation analysis, as previously described^43^, assuming the macroscopic current arises from independent, identical channels with a low probability of opening, according probability theory; *i* = σ^2^/[*I*(1 - *p*)] where *i* is unitary current, σ^2^ is the variance, *I* is mean current amplitude, and *p* is probability of opening.

### Vector construction

#### gRNA identification

CRISPR SpCas9 gRNA target sites were identified in the mouse Grid1 gene (NC_000080.6) using CRISPOR^44^. The seed sequence (GAACCCTAGCCCTGACGGCG) was chosen based on its relatively high specificity scores and the observation that it contains a Bgl I restriction enzyme site (GCCNNNN^NGGC) that overlaps with the Cas9 cleavage site.

#### mGrid1 genotyping

C57BL/6J mouse genomic DNA was isolated from tail biopsies or brain pieces containing microdissected DR by digestion in DNA lysis buffer (50 mM KCl, 50 mM Tris-HCl (pH 8.0), 2.5 mM EDTA, 0.45% NP-40, 0.45% Tween-20, 0.5 ug/mL proteinase K) for 3 h at 55 °C, and 1 h at 65 °C. Lysates were then used as templates to amplify a 654 basepair fragment including the 390F gRNA target site using Q5 HotStart Master mix (New England Biolabs). A portion of the finished PCR reaction was treated with Bgl I restriction enzyme (New England Biolabs) for 60 minutes and processed on an AATI fragment analyzer.

#### Construction and packaging of AAV vectors

The AAV vector plasmid encoding SpCas9^45^ (pX551) expressed from the MeCP2 promoter was a gift from Feng Zhang (Addgene plasmid # 60957, AAV-Cas9). The AAV packaging plasmid encoding a nuclear envelope-embedded eGFP reporter (Addgene 131682) was constructed by amplifying the KASH domain from (Addgene 60231, a gift of Feng Zhang) and fusing it (in-frame) to the end of coding region for eGFP in (Addgene 60058, pOTTC407) using ligation-independent cloning (AAV-empty, Fig. S8). gRNA was cloned into a mU6 expression cassette and then moved into an AAV backbone expressing a nuclear envelope-embedded (KASH-tagged) eGFP reporter (Addgene 131683) by PCR amplification and ligation-independent cloning (AAV-GRID1). Insert-containing clones were verified by sequencing and restriction fragment analysis prior to virus production. All AAV vectors were produced using triple transfection method as previously described^46^. All vectors were produced using serotype 1 capsid proteins and titered by droplet digital PCR.

#### Stereotaxic intracranial microinjections

Mice were anesthetized with a cocktail of ketamine/xylazine, immobilized in a stereotaxic frame (David Kopf Instruments), and received one midline injection of a 1:1 (v/v) cocktail of viruses AAV-Cas9 and AAV-empty or AAV-GRID1 for total volume 400 nL delivered over 4 minutes. The coordinates for injection were AP −4.4; ML 1.19, 20° angle; DV −3.62 mm, with respect to bregma. Prior to surgery, mice were injected subcutaneously with warm saline (0.5 mL) to replace fluid lost during surgery and given carprofen (5 mg/kg) post-surgery for pain relief. Mice recovered for >4 weeks to allow expression. Infected neurons were identified in the slice by visualization of eGFP.

### Materials

MK-801, noradrenaline, NASPM, NBQX, WAY-100635, and picrotoxin were purchased from Tocris Bioscience. All other chemicals were obtained from Sigma-Aldrich.

### Data analysis and visualization

Data were analyzed using Clampfit 10.7. Data are presented as representative traces, or in scatter plots where each point is an individual cell, and bar graphs with means ± SEM. Unless otherwise noted, *n* = number of cells. E_rev_s were determined by linear regression for each cell. Recordings in which current did not cross 0 pA were omitted from analysis. To minimize space-clamp errors, analysis of current during voltage ramps was limited to −10 mV where the currents were typically less than 500 pA. Ramp currents were averaged in 2 mV bins (20 ms). Data sets with n>30 were tested for normality with a Shapiro-Wilk test. When possible (within-group comparisons), significant differences were determined for two group comparisons by paired t-tests, Wilcoxon matched-pairs signed rank test, and in more than two group comparisons by nonparametric repeated-measures ANOVA (Friedman test). Significant mean differences in between-group comparisons were determined for two group comparisons by Mann Whitney tests, and in more than two group comparisons by Kruskal-Wallis tests. ANOVAs were followed, when p<0.05 by Dunn’s multiple comparisons post hoc test. Linear trends were analyzed using a mixed model ANOVA. A difference of p<0.05 was considered significant. Exact values are reported unless p<0.0001 or >0.999. Statistical analysis was performed using GraphPad Prism 8 (GraphPad Software, Inc.).

## ACKNOWLEDGMENTS

This work was supported by the Intramural Research Program at the National Institute on Drug Abuse (S.C.G., H.S.H, K.M.) and the National Institutes of Health Center on Compulsive Behaviors (S.C.G.). The selection and functional validation of the Grid1 gRNA and the construction and packaging of AAV viral vectors, were performed by the Genetic Engineering and Viral Vector Core of the National Institute on Drug Abuse. The opinions expressed in this article are the authors’ own and do not reflect the views of the NIH/DHHS. We thank Drs. John T. Williams and Bruce P. Bean for comments on this manuscript.

## AUTHOR CONTRIBUTIONS

S.C.G. designed the electrophysiology experiments. S.C.G. and H.S.H. performed the experiments. S.C.G. and K.M. analyzed the data and prepared the figures. S.C.G. and K.M. wrote the manuscript with the help of all co-authors.

---

**Figure S1.**
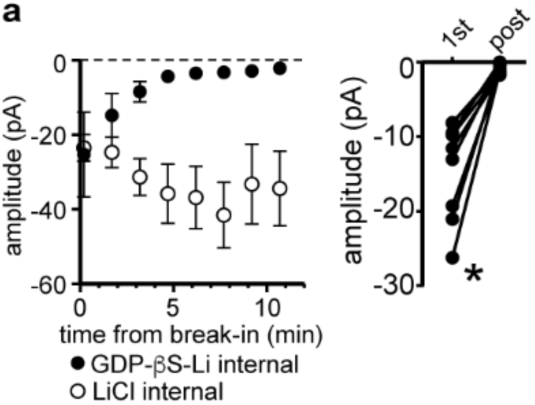
The α1-A_R_-EPSC requires G protein signaling. **a**, With GDP-βS-Li-containing internal solution, the amplitude of the α1-A_R_-EPSC ran down within ~5-10 minutes of break-in to whole-cell mode; shown in a plot compared with control internal solution containing LiCl (left) and in grouped data (right, p=0.004, n=9, 1^st^: first EPSC; post: post-dialysis). Line and error bars represent mean±SEM, * denotes statistical significance.

**Figure S2.**
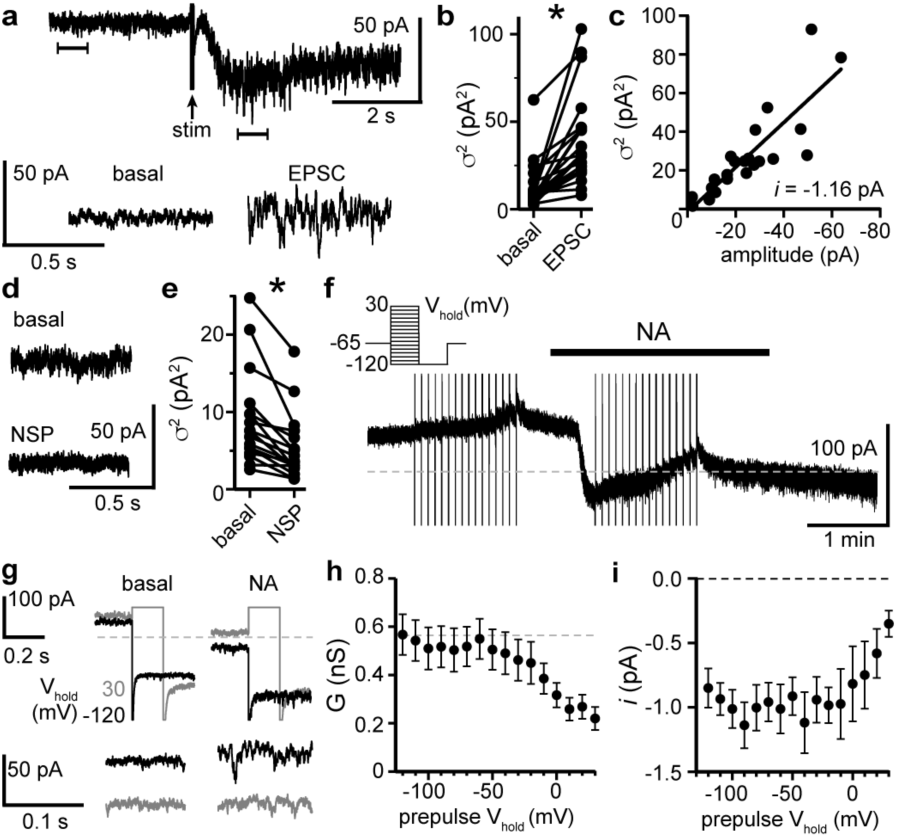
Fluctuation analysis and voltage-dependence of α1-adrenergic receptor-dependent inward current. **a**, Representative trace of membrane noise during the α1-A_R_-EPSC, brackets denote segments shown below on an expanded scale. **b**, Membrane noise (variance, σ2) increased during the α1-A_R_-EPSC (p<0.0001, n=22). **c**, Plot of α1-A_R_-EPSC variance versus mean amplitude, linear fit represents mean unitary current (*i*, r^2^=0.713). **d**, Representative trace of membrane noise under basal conditions and after NASPM application (NSP, 100 μM). **e**, Membrane noise (variance, σ^2^) decreased following NASPM (NSP, p<0.0001, n=19). **f**, I_NA_ was isolated using a two-pulse voltage protocol by subtracting current measured during voltage steps (shown in left inset) at the peak of inward current from NA application from current measured in control conditions just prior to NA (30 μM) application. **g**, Representative traces of current at V_hold_ −120 mV following 150 ms conditioning pre-pulses to −120 mV (black) or to 30 mV (grey) in control conditions (basal) and after NA. Traces below shown membrane noise of current at V_hold_ −120 mV on an expanded scale. **h**, Conductance (G_NA_) was measured at peak I_NA_ from V_hold_ −120 mV, calculated using a E_rev_ of −25.1 mV. Plot of G_NA_ versus V_hold_ of the conditioning pre-pulses, demonstrating decreased G_NA_ with depolarizing pre-pulses (linear trend of −0.022 nS/mV, p<0.0001). **i**, Plot of unitary current (*i*) measured at V_hold_ −120 mV versus V_hold_ of the conditioning pre-pulses demonstrating a decrease in *i* with depolarizing pre-pulses (linear trend of 0.026 pA/mV, p=0.002). Line and error bars represent mean±SEM, * denotes statistical significance.

**Figure S3.**
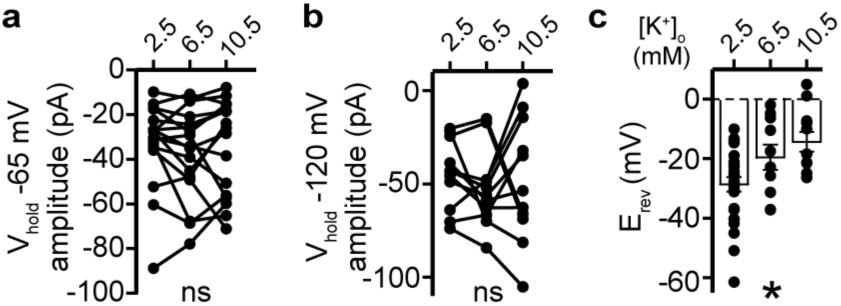
The α1-A_R_-EPSC is not due to closure of potassium conductance. **a**, Plot of α1-A_R_-EPSC amplitudes measured at V_hold_ −65 mV, in 2.5, 6.5, and 10.5 mM [K^+^]_o_ (p=0.162, n=17). **c**, Plot of α1-A_R_-EPSC amplitudes measured at V_hold_ −120 mV, in 2.5, 6.5, and 10.5 mM [K^+^]_o_ (p=0.692, n=11). **d**, Plot of α1-A_R_-EPSC E_rev_s with varying concentration of external K^+^ ([K^+^]_o_), demonstrating a depolarizing shift in E_rev_ as external K^+^ was increased (p=0.010, n=26, 10,&11). Line and error bars represent mean±SEM, * denotes statistical significance, ns denotes not significant.

**Figure S4.**
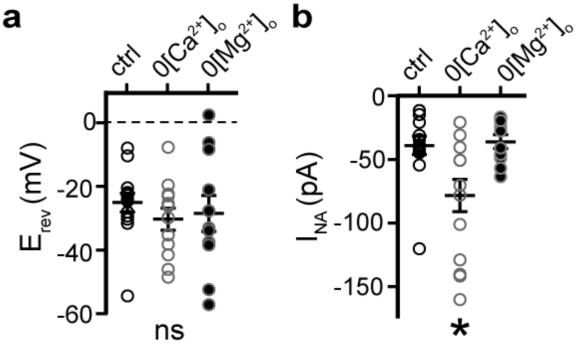
Effects of external divalents on inward current produced by noradrenaline. **a**, Plot of reversal potentials (E_rev_) of I_NA_, demonstrating no significant difference between control conditions (con), or after removal of external Ca^2+^ (0[Ca^2+^]_o_) or Mg^2+^ (0[Mg^2+^]_o_). **b**, Plot of the amplitude of I_NA_ (V_hold_ −65 mV) demonstrating an augmented I_NA_ amplitude in 0[Ca^2+^]_o_ (p=0.017, n=14), but not in 0[Mg^2+^]_o_, (p>0.9999, n=10) as compared with control conditions (n=14). Line and error bars/shaded area represent mean±SEM, * denotes statistical significance, ns denotes not significant.

**Figure S5.**
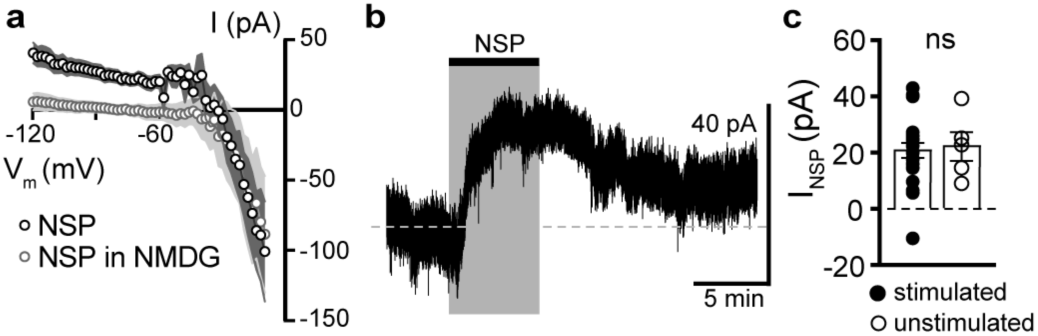
NASPM-sensitive tonic Na^+^ current does not require stimulation of the brain slice. **a**, Current-voltage relationship of the apparent outward current produced by NASPM (NSP, 100 μM). Replacing 126 mM NaCl with NMDG eliminated the apparent outward current, suggesting a block of a tonic inward Na^+^ current. **b**, Representative trace of whole-cell voltage-clamp recording of the apparent outward current induced by application of NASPM in an unstimulated brain slice. Dashed line is at 0 pA (V_hold_ −65 mV). **c**, Plot of amplitude of NASPM-induced apparent outward current in stimulated and unstimulated brain slices demonstrating no effect of prior electrical stimulation (p=0.850, n=21&5). Line and error bars/shaded area represent mean±SEM, ns indicates not significant.

**Figure S6.**
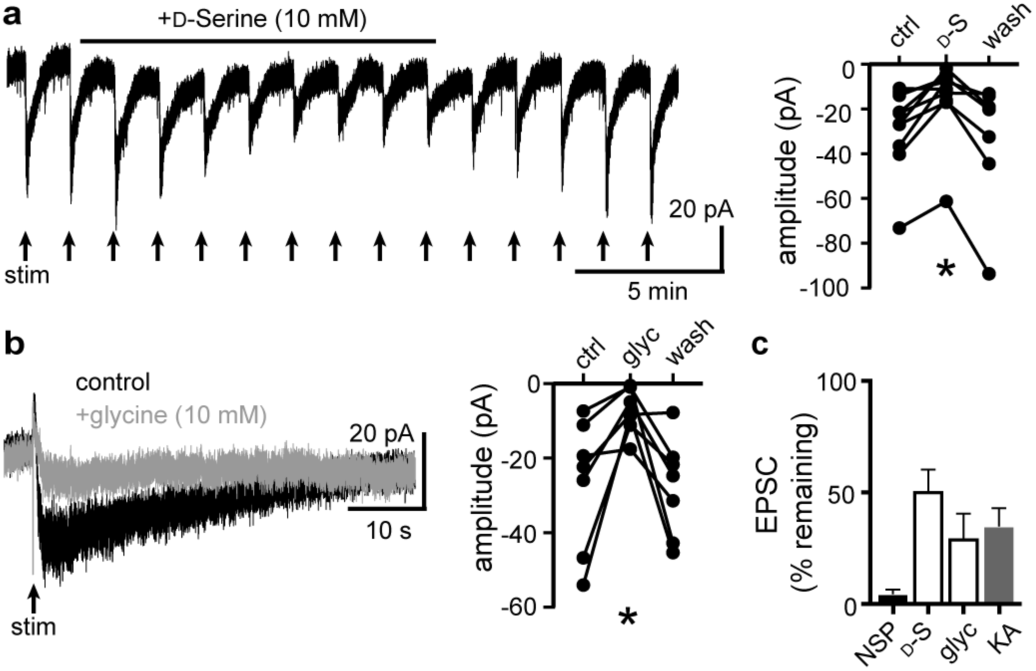
D-serine and glycine reduce the α1-A_R_-EPSC. **a**, Application of D-serine (10 mM) reversibly reduced the α1-A_R_-EPSC, shown in a representative trace (left) and in grouped data (right, p=0.001, n=7). **b**, Application of glycine (10 mM) reversibly reduced the α1-A_R_-EPSC, shown in representative traces (left, baseline adjusted) and in grouped data (right, p=0.015, n=7). **c**, Summarized data of percent remaining of the α1-A_R_-EPSC amplitude after NASPM (NSP, 100 μM), _D_-serine (_D_-S, 10 mM), glycine (glyc, 10 mM), or kynurenic acid (KA, 1 mM). Line and error bars represent mean±SEM, * denotes statistical significance, ns denotes not significant.

**Figure S7.**
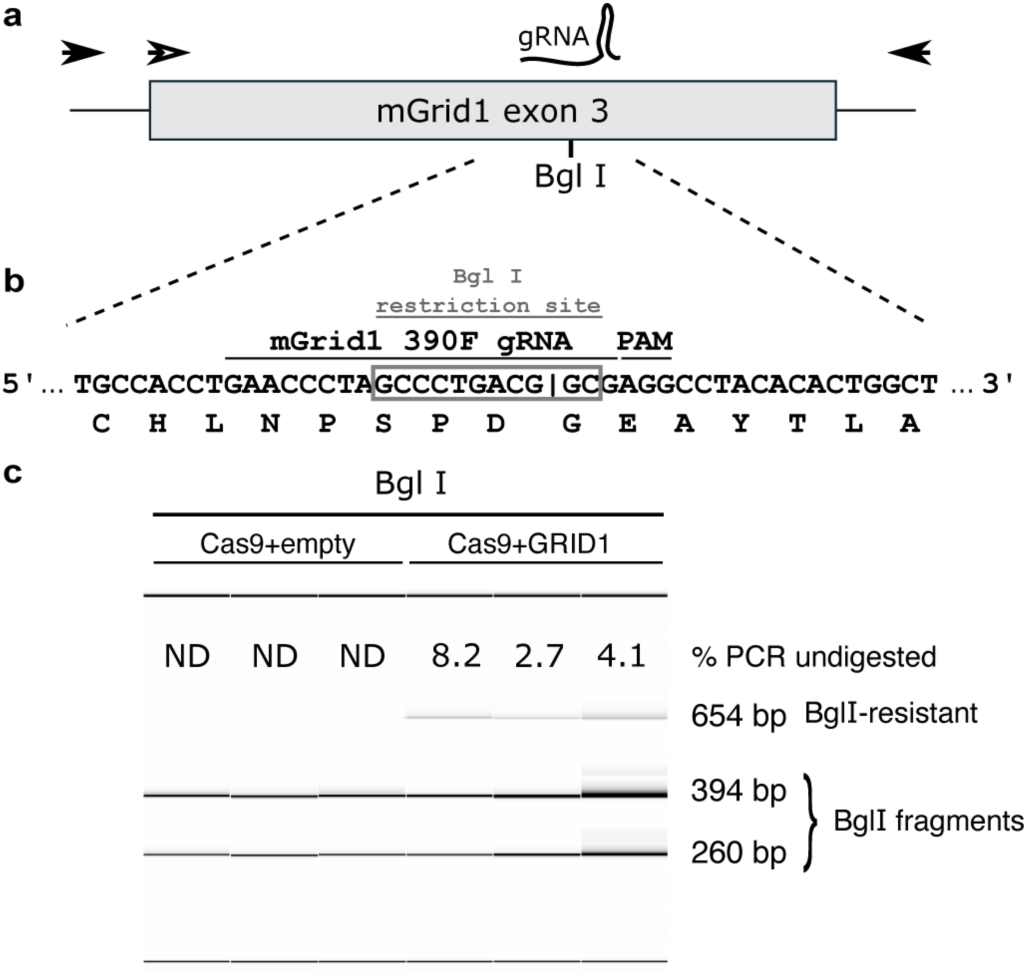
Design and testing of guide RNA targeting mGrid1. **a**, A graphic representation of the third exon of mGrid1, marked with the location of the gRNA (390F) and the overlapping BglI restriction site. Filled arrows represent the location of primers used for PCR amplification while the hollow arrow indicates the primer used for sequencing (Table S1). **b**, A nucleotide representation of gRNA target site with the translated peptide sequence underneath. The BglI recognition site (GCCNNNN^NGGC) is contained by the grey box and overlaps with the predicted Cas9 cleavage site (|). **c**, Fragment analysis on BglI-digested Grid1 PCR products amplified from the DR of mice injected with (AAV-Cas9 and AAV-empty) or (AAV-Cas9 and AAV-GRID1). “% PCR undigested” refers to the amount of the undigested PCR product relative to the total PCR product as determined by AATI fragment analysis.

**Figure S8.**
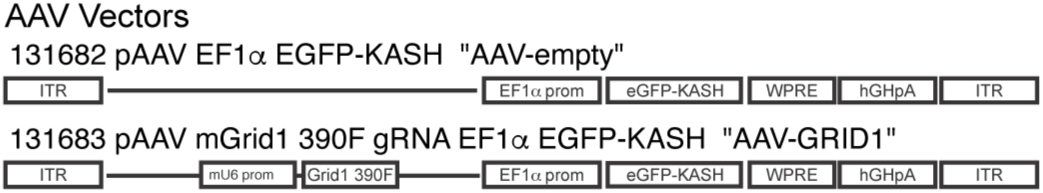
AAV vectors. A graphic representation of AAV vectors used in conjunction with AAV-Cas9 to encode eGFP reporter only (top, AAV-empty) or mouse Grid1 guide RNA with eGFP reporter (bottom, AAV-GRID1).

**Table S1.**
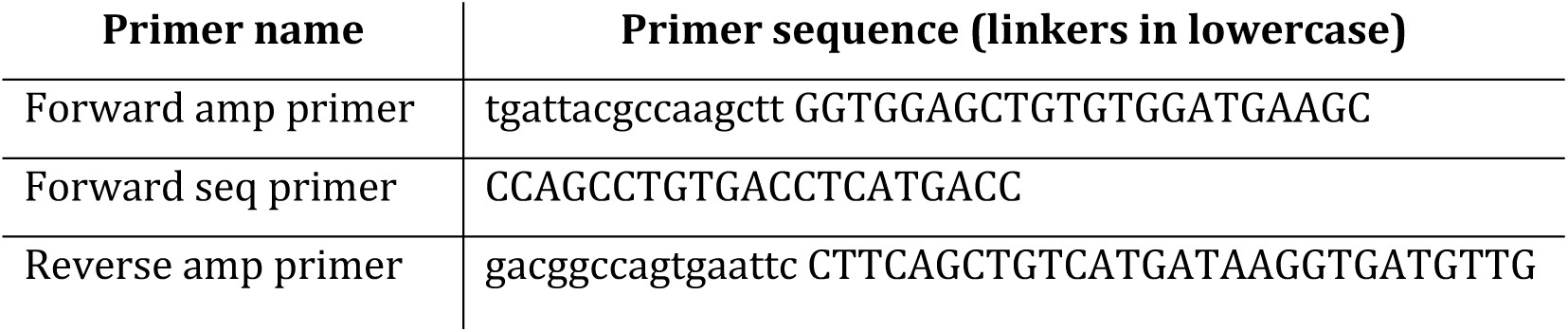
Primer sequences used in PCR amplification (amp) and sequencing (seq).

